# Loss of a belowground mutualist disrupts a fundamental biodiversity-disease relationship

**DOI:** 10.64898/2026.09.23.753874

**Authors:** Nichole A. Laggan, Aram S. Kübler, Sören E. Weber, Pascal A. Niklaus, Fletcher W. Halliday

**Affiliations:** Department of Botany and Plant Pathology, Oregon State University, Corvallis, Oregon, USA 97331; Department of Evolutionary Biology and Environmental Studies, University of Zurich, 8057, Zurich, CH; Department of Biology, West Virginia University, Morgantown, West Virginia, USA 26506

**Author notes:** Correspondence: 2701 SW Campus Way, Corvallis, OR 97331, 2701 SW Campus Way, Corvallis, OR 97331. Co-first author (Nichole A. Laggan, Aram S. Kübler).

**Keywords:** biodiversity, disease, dilution effect, host competence, mutualism, mycorrhizae, host-symbiont interactions

## Abstract

Diverse ecological communities can reduce pathogen transmission; a phenomenon known as the dilution effect of biodiversity. This effect, in turn, depends on variation in host species’ ability to spread disease (i.e., host competence) and which host species persist under biodiversity loss. Mutualistic symbionts can influence host competence and are sensitive to variation in host biodiversity; yet whether mutualists can modify dilution effects is unknown, partially because quantifying host competence during epidemics remains challenging. We experimentally tested whether mutualists could modify dilution effects by establishing gradients of host species richness, community competence, and access to the below-ground arbuscular mycorrhizal fungus (AMF), *Rhizoglomus irregulare*. Plant hosts were then exposed to the fungal root pathogen, *Rhizoctonia solani*, and disease spread was monitored over the course of an epidemic. Overall, host richness did not affect epidemic outcomes. Instead, AMF inoculation increased disease risk via species-specific increases in host competence. Moreover, communities containing highly competent host species experienced more disease, but only in the presence of mutualistic AMF. These results underscore the importance of plant community composition in mycorrhiza-mediated disease dynamics and suggest that predicting disease in our changing word might require a deeper understanding of host-symbiont and pathogen-symbiont interactions under global change.

## Introduction

Global biodiversity is declining at an unprecedented rate, with important consequences for the emergence and spread of infectious diseases (Keesing et al., 2010; Mahon et al., 2024; Halliday et al., 2021). Declining biodiversity can create positive feedback, where changes in species richness have cascading effects that further alter host community characteristics. However, the extent to which the relationship between biodiversity and disease spread is driven by host community composition remains uncertain. Furthermore, host-associated mutualisms may influence disease risk and are sensitive to the same global change drivers that cause host biodiversity loss (Weber et al., 2019; Fernandez et al., 2023). The degree to which mutualists influence disease under biodiversity loss is largely unknown. Here we employed a tractable model system using European meadow species in experimental mesocosms to explore how different host plant community compositions varying in host plant species richness and access to a below-ground mutualist (arbuscular mycorrhizal fungi) influence disease dynamics.

Highly diverse ecological communities often experience less disease transmission than communities with lower diversity (Civitello et al., 2015; Halliday & Rohr, 2019; Magnusson et al., 2019; Huang et al., 2016; Liu et al., 2018). This relationship is often described as a dilution effect. However, the opposite relationship (increasing diversity results in more transmission) is also found and described as an amplification effect (e.g., Keesing et al., 2006). A growing body of research has aimed to clarify the conditions under which disease dilution and amplification occur (Johnson et al., 2015; Rohr et al., 2020; Halliday et al., 2020; Keesing et al., 2021). An important pattern emerging from this work suggests that dilution effects most commonly occur when a reduction in species richness is accompanied by an increase in host community competence (e.g., Johnson et al., 2013), as is often the case during host community disassembly following human disturbance (Johnson et al, 2015; Halliday et al., 2020; Keesing et al., 2021). This pattern is expected because the species best able to spread disease (i.e., the most competent species) tend to possess life-history characteristics that allow them to persist in the face of human disturbance (Gibb et al., 2020). Consequently, emerging studies have attributed dilution effects more to changes in host community competence than to changes in species richness, per se (Rosenthal et al., 2022; Johnson et al., 2024).

Central to predicting dilution effects in the context of host community competence is variation in host species’ ability to spread disease (i.e., host quality; Keesing & Ostfeld, 2021; Mommer et al., 2025). High-quality hosts increase disease within a community to a greater extent than low-quality hosts. In other words, high-quality hosts are often highly competent hosts (i.e., Stewart-Merrill et al., 2020; Mommer et al., 2025). Dilution effects are expected when species in the community differ in their capacity to transmit infection when they are exposed to a pathogen (i.e., differ in host quality/competence) and when species that have a high capacity to transmit the pathogen (i.e., highly competent, high-quality, or so-called amplifying species) persist as biodiversity is lost (Johnson et al., 2015; Stewart Merrill & Johnson, 2020). Hereafter, we use the term “dilution effects” to describe situations in which an increase in host species richness involves adding low-quality hosts rather than high-quality hosts (Johnson et al., 2013; Keesing et al., 2021).

Scaling from the quality of individual host species to host community competence has become a central feature in research on dilution effects (Stewart Merrill et al., 2020, 2022; Johnson et al., 2024; Keesing et al., 2021), but quantifying species-level contributions remains challenging. Host competence is dynamic and complex and requires information on pathogen entry and persistence, host survival to transmission, and how infection outcomes vary over natural exposure gradients (Stewart Merrill et al., 2022; Stewart Merrill & Johnson, 2020). Quantifying these factors is feasible in a lab environment for some host-pathogen combinations (e.g., Stewart-Merrill et al., 2022), but whether these measurements represent actual host competence during biodiversity loss remains uncertain. For example, the loss of a competitor could increase competence of the remaining hosts by increasing host nutrition, such that low-competence host species in diverse communities become high-competence host species in species-poor communities. It may therefore be important to quantify the competence of hosts in the context of whole host communities rather than on isolated hosts. In this situation, laboratory measurements of host competence might not adequately capture host community competence under biodiversity loss.

Host competence highlights the relationship between pathogens and their hosts, but the interactions of other symbiotic organisms within hosts (e.g., mutualists) may influence host-pathogen relationships. This suggests that other (non-pathogenic) host-symbiont interactions may contribute to dilution effects and biodiversity decline. Symbionts associated with hosts can influence disease risk in various ways (Bass et al., 2019). For instance, symbionts can play a significant role in protecting their hosts from pathogens in multiple plant and animal systems (Bass et al., 2019; Haine, 2008), but even these so-called “defensive mutualists” can increase host susceptibility in some cases (Halliday et al., 2017). Anthropogenic disturbances have the potential to cause local extinction of mutualists and disrupt mutualistic relationships, presenting a global risk with significant consequences for biodiversity loss (Aslan et al., 2013; Toby Kiers et al., 2010).

In plant hosts, arbuscular mycorrhizal fungi (AMF) might be particularly important symbionts for influencing disease outcomes and host competence. AMF (Glomeromycota) are associated with more than 80% of current land plants, widely considered the most widespread fungal symbionts of plants (Bonfante & Genre, 2010; Smith & Read, 2008), and ongoing global changes are expected to not only reshape the composition of AMF but also redefine the nature of their mutualistic interactions with plants (Weber et al., 2019). At the same time, the impact of AMF on plant fitness and their role in pathogen protection are widely acknowledged (Bender et al., 2019; Jung et al., 2012). AMF can improve plant fitness by influencing soil nutrient transfer, increasing biomass, and occasionally improving resistance to abiotic stress and pathogens (Bonfante & Genre, 2010; Smith & Read, 2008). The presence of AMF in the soil can induce changes in the rhizosphere environment, including changes in the physical and chemical properties of soil, enhancing the growth of beneficial microorganisms and competition with pathogenic microorganisms (Weng et al., 2022).

Impacts of AMF on host traits (including host competence) often vary across host functional groups (Romero et al., 2023), species (Kuyper & Jansa, 2023), or even genotypes (Eck et al., 2022), suggesting that AMF could alter host community competence, and thereby impact whether and when dilution effects are observed. Not all plants benefit from AMF (Kuyper & Jansa, 2023), and in some cases, associating with AMF does not provide advantages to a host plant (Holland et al., 2019). A field study found that AMF inoculation increased host plant growth but also increased pathogen infection rate, while providing some protective effects to host genotypes that were more susceptible to pathogen infection (Eck et al., 2022). Together, these studies suggest that host competence (and thus dilution effects) could depend strongly on AMF, and that these effects might vary across host taxa.

AMF-modification of dilution effects might be most relevant in the context of soil-borne pathogens. Indeed, soil-borne pathogens, such as fungi, pose major threats to plant health (Raaijmakers et al., 2009). In these systems, the belowground transmission of fungal pathogens occurs when hyphal growth extends from the roots of an infectious plant and reaches the roots of a susceptible plant (Stacey et al., 2001). Heterogeneous transmission patterns of soil-borne fungal pathogens can occur in mixed populations with varying infectivity and susceptibility among plant species (Otten et al., 2005). The formation of a parasitic relationship between the plant host and the pathogen occurs in the rhizosphere, where interactions with the rhizosphere community of microorganisms (including AMF) can shape the outcome of pathogen infection (Raaijmakers et al., 2009). For example, AMF could amplify soil-borne disease transmission by increasing connectivity among plant roots, or alternatively, AMF could mitigate disease by inhibiting the hyphal growth of soil-borne pathogens or pre-empting host tissue from becoming infected

There is widespread recognition that biodiversity loss and the disruption of mutualisms increase disease risk with crucial implications for infection outcomes in ecological communities and in agriculture. However, how these factors combine and interact to influence disease epidemics remains poorly understood. In this study, we aimed to elucidate the role of AMF in shaping the impact of biodiversity on disease risk in plant communities. We examined whether AMF could enhance or reduce young plant survival under exposure to a soil-borne root pathogen and were interested in understanding whether this effect might change with increasing plant species richness. To address this question, we leveraged a model system for exploring dilution effects using the ubiquitous fungal pathogen, *Rhizoctonia solani*. *R. solani* is a soil-borne pathogen, primarily of agricultural concern in seedlings, and infects near the soil line, causing seedling collapse and ultimately death of the plant (i.e., damping-off disease) (Otten et al., 2003; Rosenthal et al., 2022). However, in addition to damping-off disease, symptoms on diverse host plant species can also include seed rot, root rot, hypocotyl rot, crown rot, stem rot, limb rot, pod rot, stem canker, black scurf, and seedling blight (Ajayi-Oyetunde & Bradley, 2018). The effective transmission of the pathogen, which gradually decelerates over time, depends on the critical factors of the distances between potential neighboring host plants and the availability of resources (Bailey et al., 2000; Rosenthal et al., 2022). This model system has been used to explore the effects of neighboring identity on disease in wild plants (Ampt et al., 2022) and to study the role of community competence and assembly mechanisms on dilution effects in cultivated, agricultural plants (Rosenthal et al., 2022).

Here, we leverage a model system of community competence and manipulate the presence and absence of a mycorrhizal mutualist, the AMF species *Rhizoglomus irregulare* (syn. *Rhizophagus irregularis).* We assess community competence between multiple species assemblages by estimating species-level competence through single-species inoculations and through mechanistic diallel analysis to determine general combining abilities. Generally, we find that host community competence can drive disease transmission, consistent with the dilution effect hypothesis, but that this effect can be mediated by mycorrhizal mutualism. In addition, we find that species-specific host competence can be estimated from multiple approaches and that phylogenetically similar species can exhibit substantial variation in competence. Combined, these results highlight the importance of mycorrhiza-mediated disease dynamics in predicting disease risk in a changing world.

## Materials and Methods

### Experimental design

Using a mesocosm experiment, we explored the relationship between herbaceous plant diversity, arbuscular mycorrhizal mutualism, and transmission of the soil-borne pathogen *Rhizoctonia solani*. Specifically, we created plant communities consisting of one, two, three or four species (Figure 1A). We replicated this plant diversity gradient using two non-overlapping sets of four European grassland plant species, with all 15 possible plant species combinations realized within each pool. Species were evenly assigned to pools so that each pool contained one grass, one non-aster dicot, one aster, and one legume species, and so that each pool had approximately equal community-level competence.

**Figure 1.**
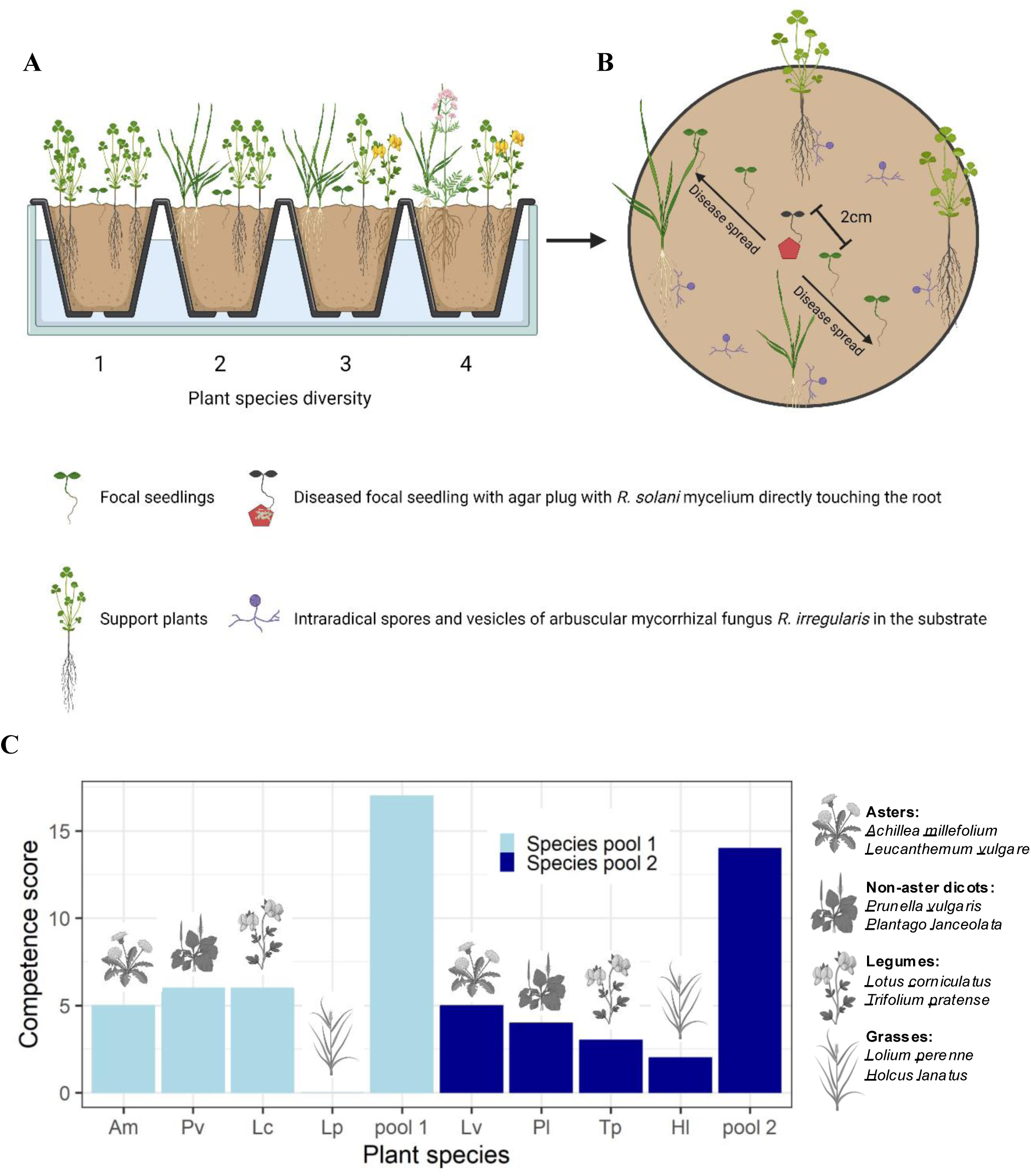
Overview of experimental design and host species competence. A) Experimental mesocosms consisted of mature support plants that were established to form a plant species diversity gradient and focal seedlings that were used to measure disease transmission. B) A detailed view of one experimental mesocosm. Support plants were placed on the border of each community with the goal of establishing a biodiversity gradient and mycorrhizal network (in the treatment where intraradical spores of arbuscular mycorrhiza of *Rhizoglomus irregularis* were added to the substrate). The five focal seedlings (*Achillea millefolium* in Species pool one and *Leucanthemum vulgare* in species pool two) were established with a two-centimeter interplant distance. Each central focal seedling was inoculated with an agar plug containing *Rhizoctonia solani* mycelium (or a sterile control plug) directly touching the root (shown as a red pentagon). C) Estimated competence of each plant species (detailed in Appendix I). Plants were arranged into species pools so that the four-species mixture had a similar “community competence” (i.e., average competence across all individuals in the pool). Figure created in BioRender.com.

We replicated this plant diversity treatment with and without inoculation with the AMF species *Rhizoglomus irregulare* (syn. *Rhizophagus irregularis,* strain QS81 from INOQ Gmbh, Germany). *R. irregulare* was selected because it is widely distributed (Błaszkowski et al., 2008) and economically important as it is part of key components in commercially available AMF products, as well as a scientifically important model organism (Sieverding et al., 2015).

To separate the effect of density of each particular host species from the effect of the plant species diversity of the community in which this host occurs, we planted a row of five seedlings of a focal host species in each plant community (Figure 1B). In each mesocosm, an epidemic was initiated by inoculating the central seedling with the fungal pathogen, *Rhizoctonia solani*. These experimental treatments were replicated three times. An additional 20 communities (monocultures and four-species mixtures with and without mycorrhizae) were planted as mock controls and never inoculated with the fungal pathogen. Overall, the experiment consisted of 200 experimental mesocosms (two plant species pools x 15 plant species compositions per pool x two mycorrhiza treatments x three replicates plus 20 non-infected controls).

### Experimental setup

Specific details of pathogen propagation and maintenance, substrate preparation and mycorrhizal inoculation, seed preparation and germination, planting of focal seedlings, and greenhouse conditions are provided in Appendix I. Briefly, potting soil (ED73 lawn soil (Einheitserde, Germany)) and sand were sterilized by high-energy X-rays (25-60 kGy,), and autoclaving, respectively, then mixed at a 2:1 ratio. The substrate was then re-inoculated with an AMF-free bacterial/rhizobia mix from an extensively managed meadow following Weber et al. (2024). Mycorrhizal inoculum was added to half of the soil at a rate of 100 mg of *R. irregulare* mycorrhizal root powder / L of soil. Soil was packed into 10 cm diameter, 500 mL, round pots, which served as individual experimental mesocosms.

Plant seeds were surface-sterilized, germinated in Petri dishes on sterilized sand, and then transplanted after five days. Planting occurred in two phases; first, the biodiversity gradient was established by planting four plants in the edges of a square in the pot (Figure 1B). These plant communities were then grown for three weeks to support the establishment of a mycorrhizal network. Next, a row of seedlings of one plant species per species pool (*Achillea millefolium* in pool 1 and *Leucanthemum vulgare* in pool two) was planted into each community to serve as focal seedlings for measuring disease transmission. Both focal species belong to the same plant family (Asteraceae) and were selected based on similar (intermediate) competence from pilot experiments (Table S1). Two days after planting, the central seedling in each pot was inoculated with *R. solani* (AG4-HGI, strain CBS 124594 from the Westerdijk Fungal Biodiversity Institute) (Figure 1B). An agar plug with actively growing mycelium (1cm^2^) was cut out of agar culture plates and placed with the mycelium side directly adhering to the primary root of the seedling and then covered loosely with soil. For the mock-inoculated control mesocosms, we used a sterile agar plug. The central seedling was selected to avoid potential edge effects from specific plant species in the plant community.

### Disease symptoms

Disease symptoms were recorded on each of the 1000 seedlings every one to two days for 20 days following inoculation of the central host individual, resulting in a total of 12 disease surveys (n = 12,000 total observations). Disease symptoms were scored above ground and included three categories: healthy, diseased (showing symptoms of leaf and hypocotyl rot, discoloring or spotting, wilting, and stem collapse (Ampt et al., 2022)), or dead. No disease was observed on any of the mock-inoculated control mesocosms.

### Root colonization by AMF

20 days after inoculation, root colonization by AMF was verified by harvesting all control mesocosms and separating the root systems of all plant individuals. The roots were washed, and fine root samples were collected, stained, and mycorrhizal colonization was determined by microscopy using the line intersect method (Vierheilig et al. 1998). Our microscopy image analysis revealed low-level AMF colonization in the AMF-inoculated mesocosms (Figures S1-S8). As expected, none of the AMF-free mesocosms had signs of AMF colonization.

### Quantification of disease burden

We quantified disease burden of all focal seedlings as area under the disease progress stairs (AUDPS) across all of the 12 surveys, following Ampt et al. (2022)

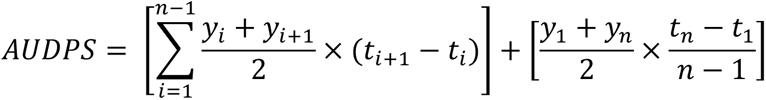

where t_i_ is the number of days after inoculation (i = 1..n), n is the total number of observations, y_i_ indicates the presence of disease (0 or 1) (Heckman et al., 2019; Simko and Piepho, 2012). Similar to the area under the disease progress curve, AUDPS integrates the development of disease progress into a single value of disease (Halliday et al., 2018; Madden et al., 2017).

### Quantification of community competence

We quantified host community competence using individual species’ host competence data from pilot experiments (Appendix I). Species competence values were used to compute a theoretical community competence for each plant community, calculated as an average of the competence of each support plant species present in the plant community.

### Statistical analysis

Eight of the total of 180 experimental mesocosms were excluded from the data analysis because they were planted with the wrong species. In one mesocosm containing four support plants, a single species was missing from the intended three-species composition, and the community’s diversity level was therefore adjusted and treated as a two-species composition (n=1). Only four of the 1,000 seedlings were excluded from data analysis because they died for reasons other than *R. solani* infection, or were damaged during surveys.

Data were analyzed using R 4.3.1 (R Core Team, 2023). To test for effects of the experimental treatments (plant species diversity, mycorrhizal inoculation, species pool, host community competence and their interactions), we used the ‘lme4’ (Bates et al., 2015) and ‘lmerTest’ (Kuznetzova et al., 2017) packages to perform linear mixed-effects models and then summarized in ANOVA tables with sequential (type I) tests.

Plant species richness (continuous), species pool (two-level factor) and AMF inoculation (two-level factor) were fixed terms, whereas plant species composition was fitted as a random term, because it is the unit of replication for testing diversity effects (Schmid et al., 2017; Schnyder et al., 2023). The combination of species composition and AMF treatment was the residual, i.e., all data were aggregated at this level prior to analysis. For model predictions and post hoc tests, we used the R packages ‘effects’ (Fox & Weisburg 2019) and ‘emmeans’ (Lenth 2023).

A total of four models were fitted using disease burden as the dependent variable. To determine the general effect of species richness and AMF inoculation on disease burden, we fitted a first model which included AMF inoculation and plant species richness as interactive fixed effects. Species composition in the plant communities was added as a random term. To investigate how the different pools respond to inoculation with AMF, we next fitted a second model identical to the previous model but added species pool as an interactive fixed effect.

To explore the effect of average community competence, we fitted a third and fourth model that were identical to the first and second models but included average community competence as interactive fixed effects in place of the plant species richness treatment. However, the interpretation of p-values and confidence intervals in these models requires caution, due to potential pseudo-replication in the x-variable. Specifically, the competence score measure for the seedling in monocultures is not error-free, and the process of generating community-competence measures from a limited set of species-specific values may introduce bias. Consequently, reported p-values and confidence intervals may be lower than actual values. To assess differences in the relationships of average community competence and disease burden between the AMF inoculation treatments, we performed post hoc tests in ‘emmeans’ to estimate slopes and statistical support.

Because host competence depends on experimental conditions, we used our disease scores to estimate the competence of the plant species that formed the community into which the seedlings were embedded. Specifically, we used mechanistic diallel analysis to model seedling disease scores as a function of additive contributions (so-called “general combining abilities”, GCA) of the adult plant species in the corners of the mesocosms. The diallel models were fitted separately for each species pool x AMF treatment combination, using data from monocultures and two-species mixtures. Models had the form y_ab_ = GCA_a_ + GCA_b_ + ε_ab_, where GCA_x_ is the general combining ability of species x (Schnyder et al., 2023). Monocultures were treated as combinations of two identical species, i.e. y_aa_ = 2GCA_a_ + ε_ab_ (Schnyder et al., 2023).

## Results

First, we tested for a general effect of species richness and inoculation with arbuscular mycorrhizal fungi (AMF) on disease burden. Overall, AMF presence increased disease burden (+17%, F_1,28_=4.332, P=0.047). However, disease burden did not depend on species richness (F_1,28_=2.33; P=0.138), nor did such an effect depend on the presence of AMF (F_1,28_=0.401; P=0.532, n=60 plant species compositions; Figure 2a; Table S2).

**Figure 2.**
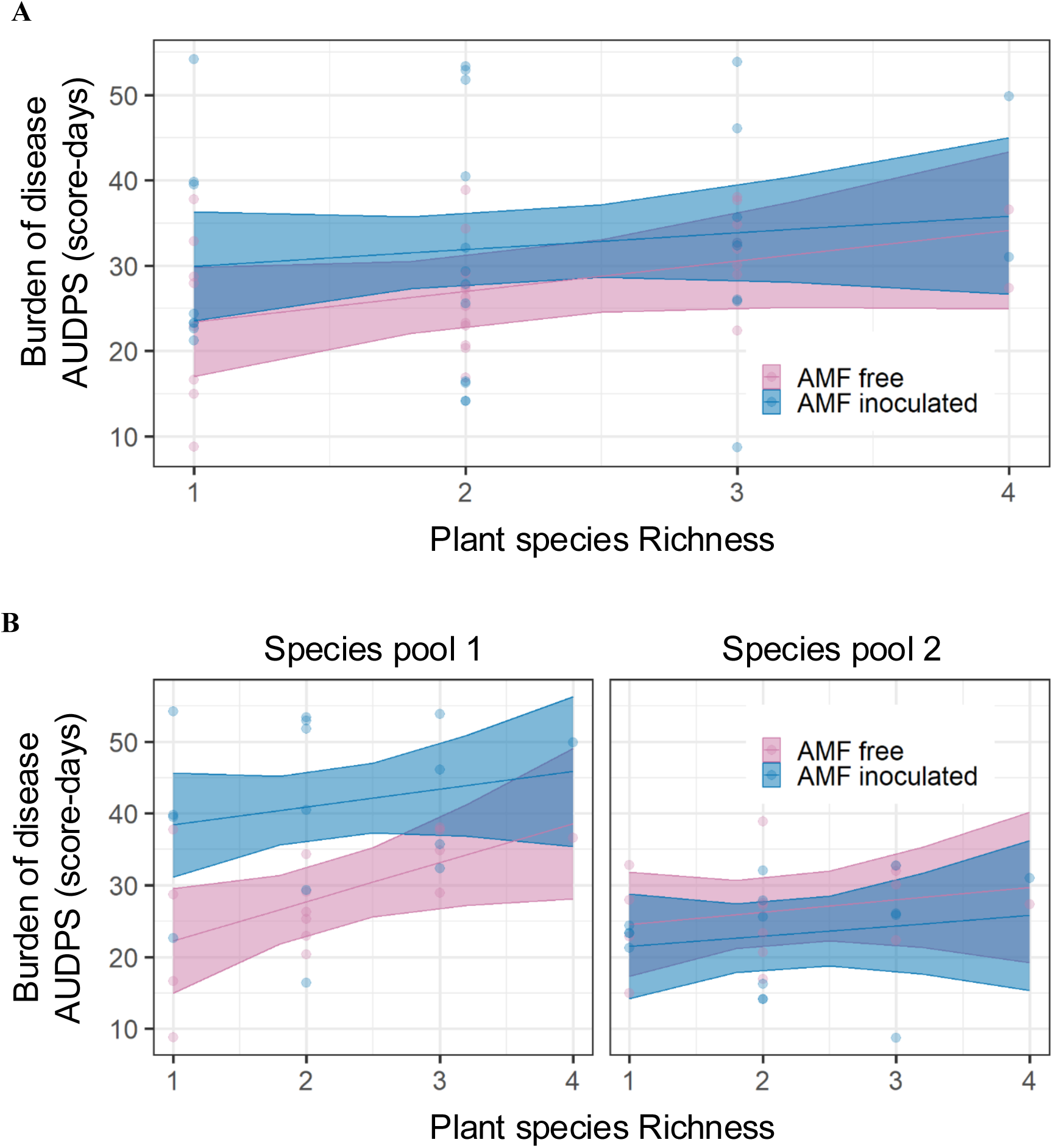
Burden of disease across the entire epidemic, measured as area under the disease progress stairs (AUDPS) and derived from the cumulative scores of 0 and 1 observed across 5 focal seedlings in 12 surveys over 20 days. A) Disease burden as a function of plant species richness and inoculation of the soil with arbuscular mycorrhizal fungi (AMF). B) Disease burden as a function of plant species richness, AMF inoculation, and plant species pool. Points represent the average value of disease burden across experimental replicates for each species composition. Lines and shaded areas represent model-estimated means and 95% confidence intervals from linear mixed-effects models. Pink shows no AMF inoculation and blue shows AMF-inoculated mesocosms. AMF inoculation increased disease burden, and this effect was generated by a strong response in Species pool one.

Next, we tested whether these effects differed between species pools. Disease burden was 42% higher in species Pool 1 than in species pool 2 (Figure 2b; Table S3; F_1,26_=14.2, P<0.001), and effects of AMF differed between pools (F_1,26_=22.610; P<0.001). Specifically, AMF inoculation increased disease burden by 42.1% in species pool 1 but not in species pool 2 (P=0.003 and 0.920 for contrasts among AMF treatments, Tukey post hoc test). Consistent with the analysis across species pools, there was weak evidence for a positive relationship between plant species richness and disease burden (Div, F_1,26_=3.4039; P=0.076). Hence, the effect of AMF was largely due to species pool one, indicating that sets of plant species with similar phylogeny can respond very differently to fungal mutualists and exhibit strongly diverging disease epidemics.

Next, we tested whether host community competence (calculated as the average competence score of the support-plant species, as quantified on seedlings in the pilot experiment) and AMF inoculation explained disease burden in focal seedlings. Across species pools, there was evidence for a statistical interaction between host community competence and AMF inoculation (Figure 3a; Table S8; Competence × Mycorrhizae, F_1,28_=7.430; P=0.011). Specifically, AMF-inoculated communities experienced more disease with increasing community competence (slope: 4.588, *t*_48.2_ = 3.273, P=0.002), but non-AMF-inoculated communities did not (slope: 0.413, *t*_48.2_ = 0.295, P=0.769). This statistical interaction between host community competence and AMF inoculation was present even when models included species pool as a fixed effect (Figure 3b; Table S9; Competence × Mycorrhizae, F_1,26_=11.631; P=0.002), despite differences between species pools in overall disease and the effects of AMF inoculation on disease. Overall, host community competence predicted disease across communities, and this prediction depended on inoculation with AMF. These results suggest that disease transmission was best explained by variation in host competence rather than host richness, manifested differently across different host species pools, and required the presence of disease-amplifying AMF in the soil.

**Figure 3.**
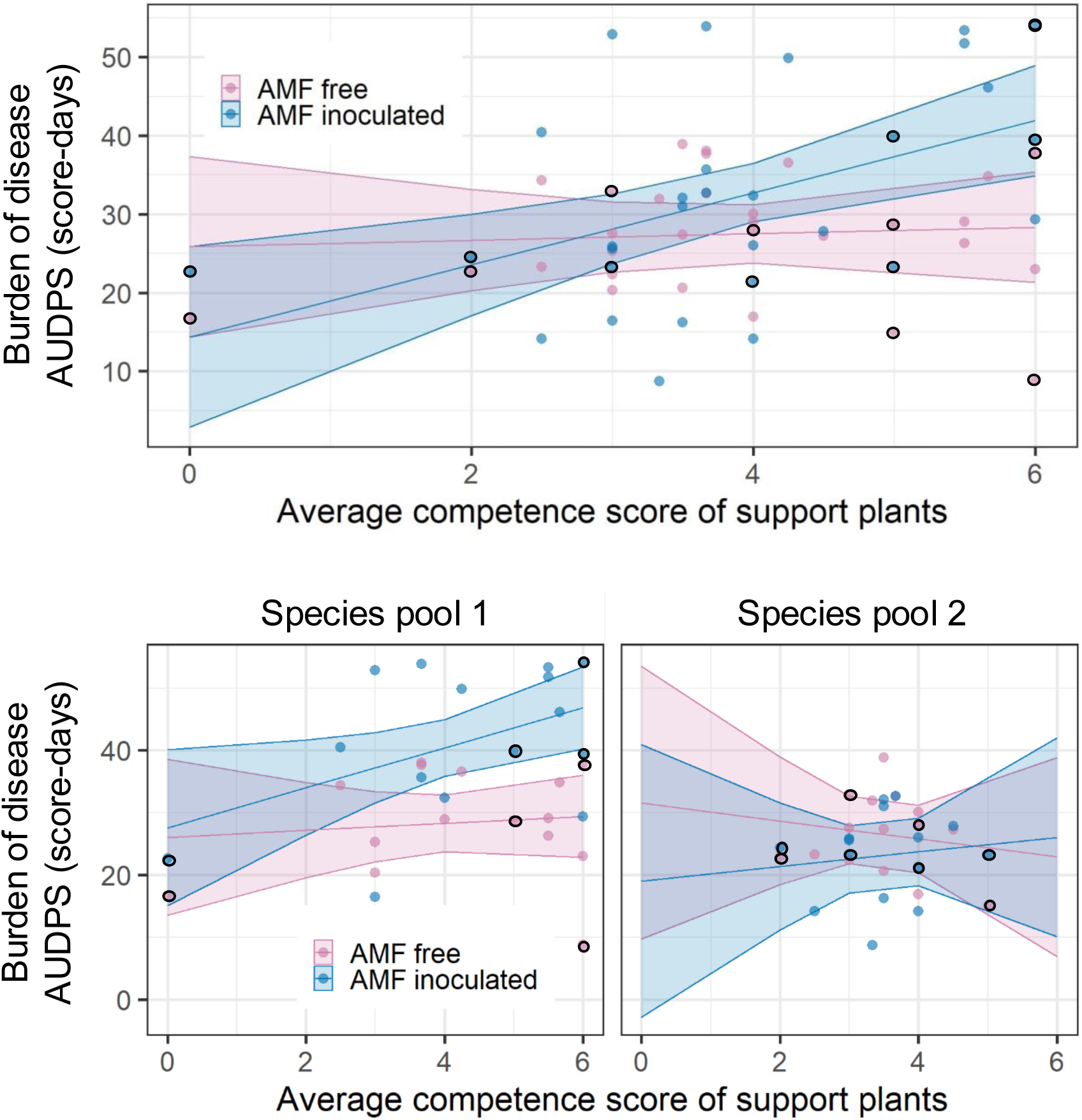
Burden of disease across the entire epidemic, measured as area under the disease progress stairs (AUDPS) and derived from the cumulative scores of 0 and 1 observed across five focal seedlings in 12 surveys over 20 days. A) Disease burden as a function of host community competence (computed as the average competence score of plant species in a community) and inoculation of the soil with arbuscular mycorrhizal fungi (AMF). B) Disease burden as a function of host community competence, AMF inoculation, and plant species pool. Points represent the average value of disease burden across experimental replicates for each species composition. Lines and shaded areas represent model-estimated means and 95% confidence intervals from linear mixed-effects models. Pink shows no AMF inoculation and blue shows AMF-inoculated mesocosms. Black circles show monocultures to highlight species-specific contributions to disease burden. Increasing host community competence increased disease burden, but only in AMF-inoculated communities; this effect was most evident in Species pool one.

We then leveraged a mechanistic diallel analysis (see Methods) to capture host competence and explore the effect of community competence on disease burden during the epidemic. In this mechanistic diallel analysis, hosts’ competence scores were estimated as general combining abilities by comparing differences in disease between monocultures and two-species mixtures. As expected, general combining abilities varied across plant species and mycorrhizal inoculation (Figure 4, panels A-D; Tables S4-S7). Moreover, species with higher competence in the pilot experiment also made greater additive contributions to disease burden in experimental mesocosms, except for *Leucanthemum vulgare* and *Lotus corniculatus,* which had very high competence scores but very low GCA estimates (Figure 4E). These results support the conclusion that individual species’ contributions to host community competence are additive and largely reflect differences in their seedling-derived competence scores.

**Figure 4.**
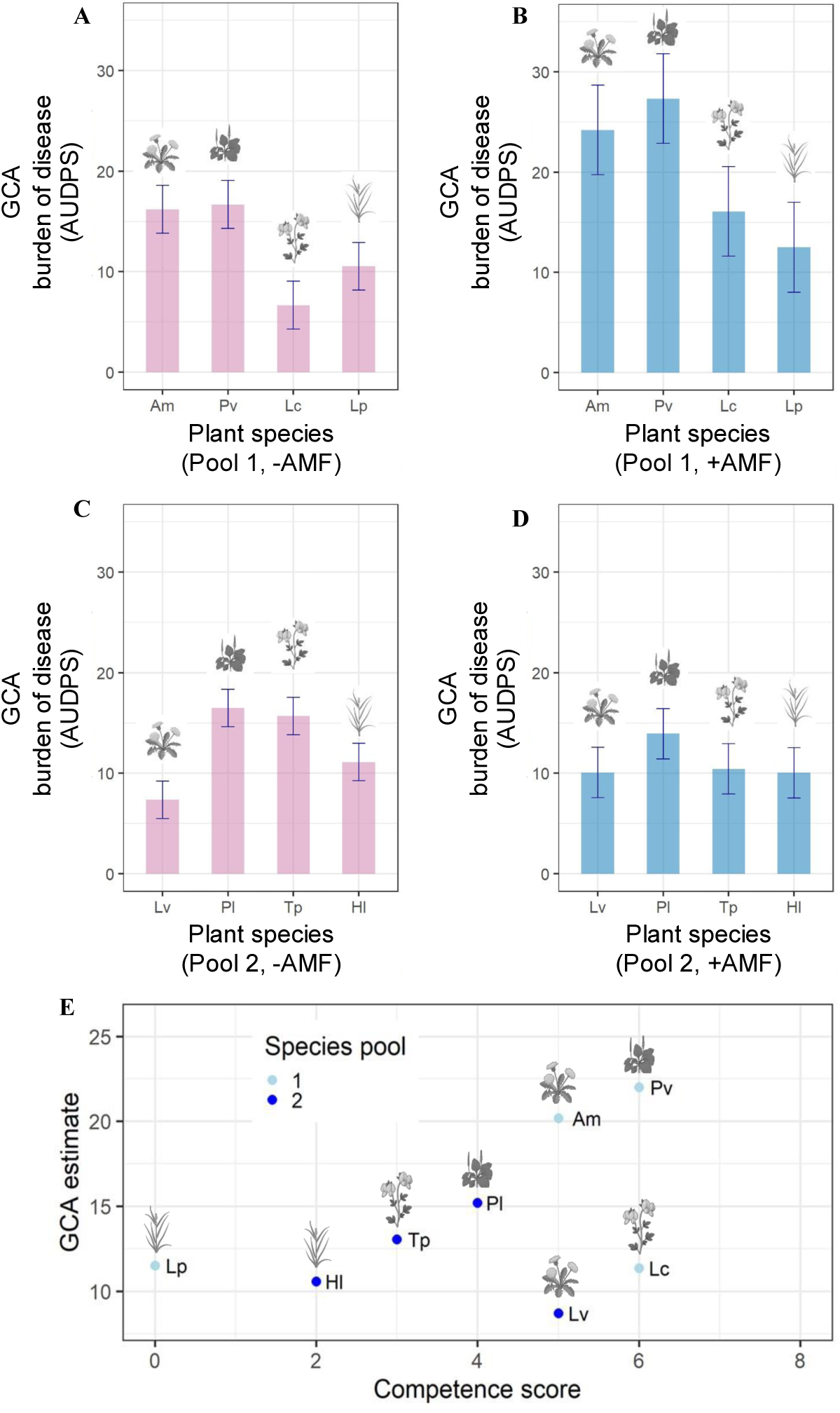
Results of mechanistic diallel analysis from single species and two-species mixtures. A-D) General combining abilities (GCAs) of plant species in two species pools with and without arbuscular mycorrhizal fungal (AMF) inoculation. GCAs indicate the additive contributions of the plant species to the disease burden of the individual compositions. Bars indicate one standard error. E) GCA estimates as a function of plant species competence scores for both plant species pools averaged across AMF treatments. Species pool 1 (Panels A and B and light blue in Panel E): Am = *Achillea millefolium,* Pv = *Prunella vulgaris*, Lc = *Lotus corniculatus*, Lp = *Lolium perenne*. Species pool 2 (Panels C and D and dark blue in Panel E): Lv = *Leucanthemum vulgare*, Pl = *Plantago lanceolata*, Tp = *Trifolium pratense*, Hl = *Holcus lanatus*. GCAs vary considerably among plant species and AMF treatment, with the highest GCAs observed in AMF-inoculated mesocosms of Species pool one (Panel B); GCAs and species competence scores are generally congruent except for *Leucanthemum vulgare* and *Lotus corniculatus*, which were highly competent but contributed little to disease transmission in experimental mesocosms.

## Discussion

While both plant diversity and belowground mutualistic microbes play key roles in ecosystems, how they interact to shape disease dynamics is poorly understood. We explored how biodiversity and mutualism interact to influence disease by conducting a mesocosm experiment using two plant species pools with an established biodiversity gradient, mycorrhizal fungi, and a fungal root pathogen. We found that increasing species richness did not influence disease, but increasing host community competence resulted in elevated disease burden. Interestingly, the loss of a belowground mutualist disrupted the disease-diversity relationship and generally reduced disease. Our study also revealed that species-specific host competence can be estimated from multiple approaches: single-species seedling inoculation experiments and mechanistic diallel analysis from infected communities. Broadly, our findings demonstrate that both host community competence and mutualistic interactions can influence disease risk during biodiversity loss.

Although previous work suggests that greater species richness may reduce disease (Civitello et al., 2015; Halliday and Rohr, 2019; Mitchell et al., 2002; Rottstock et al., 2014), we did not find clear evidence that species richness affected disease dynamics in our experimental plant communities (Figure 2A; Table S2). Instead, our results support a growing body of work that host competence is crucial for predicting disease risk (Johnson et al., 2013; Stewart Merrill et al., 2022; Stewart Merrill & Johnson, 2020) (Figure 3A, Table S8). Host competence and species richness are often coupled following anthropogenic changes to ecosystems; thus, disease ecologists have struggled to explain whether dilution effects result from changes in species richness or host community competence. Here, by experimentally decoupling host community competence (measured using mechanistic diallel analysis and species-level seedling inoculations) and species richness, we add support to the hypothesis that changes in host community competence may underlie important dilution effects as communities disassemble (Rosenthal et al., 2022; Halliday et al., 2020; Mahon et al., 2024; Jonson et al., 2015; Keesing & Ostfeld, 2021).

Dilution effects may depend on associated changes in host community competence, but measuring this is limited by researchers’ ability to collect accurate measurements of individual host species’ competence, a trait that can change across host ontogeny and in response to environmental gradients (Halliday et al., 2023; Stewart Merrill & Johnson, 2020). In our study, we leveraged a mechanistic diallel analysis as a new tool for measuring biodiversity-ecosystem functioning relationships (Schnyder et al., 2023) to quantify host competence as epidemics unfolded. This approach is appropriate only if the contributions of each species are additive and there are many species combinations, as was the case in our experimental mesocosms. We identified distinct general combining abilities (GCAs) among adult plant species, indicating variation in their impact on the overall disease burden observed in the focal seedlings (i.e., variation in neighboring host competence; Ampt et al., 2022). Further, there was an association between a species’ pre-measured seedling competence score from pilot experiments and its GCA value for most of the eight species employed (Figure 4E). This implies that, under certain conditions, laboratory measures of host competence might be sufficient to predict dilution effects. However, two species had very high competence scores, but very low GCA estimates (Figure 4E). Differences in disease transmission estimates may arise from factors such as plant age or interactions among the plant species in mixture, given that the competence score was based on younger monoculture seedlings. As indicated by previous studies, plant resistance may increase with host age, dependent upon host species identity, thereby limiting transmission by outpacing pathogen growth (Bruns et al., 2022; Otten et al., 2003). Alternatively, increased root interweaving in older plants may enhance belowground pathogen transmission (Bailey et al., 2000). This aligns with the findings by Ampt et al. (2022), which suggest that disease transmission is influenced by both successful pathogen infection and the development of resistance, a process regulated by the identity and developmental stage of neighboring and focal plants. Moreover, we found that overall disease burden differed between our two species pools (Figure 3B-C; Figure 4A-D; Table S9), despite being theoretically constructed with equal community competence, underscoring the need to consider individual species competence in the context of biodiversity decline and ongoing epidemics.

We expected AMF inoculation to influence disease dynamics through their impact on host competence; however, in contrast with the common observation that AMF reduce disease, we found that AMF inoculation increased disease burden in some of our plant communities (Figure 3A-B; Tables S8-S9). AMF are often thought to enhance plant resistance to fungal pathogens (Borowicz, 2001) and have been promoted as a biological control agent against pathogen-induced wilt and for influencing plant secondary metabolites in agricultural cultivars (Hao et al., 2005; Hu et al., 2010). However, conflicting evidence from greenhouse studies (Holland et al., 2019) and observations in natural ecological conditions reveals the complexity of mycorrhizal fungi’s benefits and risks to wild host plants, including increased infection rates (Eck et al., 2022). Our findings suggest that species interactions that are commonly considered mutually beneficial can have unexpected effects on infectious disease dynamics.

Despite a measurable effect of AMF inoculation on disease dynamics, AMF inoculation did not always result in consistent AMF colonization in this experiment. In general, wild plants are often more sensitive to variations in mutualistic AMF than domesticated agricultural plants, potentially resulting in reduced mutualistic effects in commercial AMF isolates (Kokkoris et al., 2019). Further, relatively benign experimental conditions could have reduced plants’ need to associate with AMF (Holland et al., 2018). Plants often experience reduced root colonization when soil fertility is high, indicating that fungal symbionts may not provide benefits to plants under these circumstances (Smith and Read, 2008). In contrast with past studies that fertilized mesocosms with a low-phosphorus nutrient solution (Romero et al., 2023; Van Der Heijden et al., 1998), we did not fertilize the mesocosms in this experiment. The absence of fertilizer may explain the notably low AMF colonization in our experimental root samples, highlighting a limitation in our study. Regardless, we did observe an effect of AMF inoculation on disease burden, which suggests that even low-level AMF colonization rates and the presence of AMF in the soil can influence pathogen abundance.

To our knowledge, this is the first study to show that AMF inoculation can cause species-specific enhancement of disease risk in ecological communities. This result is consistent with past studies showing host-specific responses to AMF in different contexts (e.g., growth or Phosphorus acquisition; Van Der Heijden et al, 2015; Romero et al., 2023). This species-specific influence of AMF on disease was further supported by the mechanistic diallel analysis revealing that individual plant species contributed differently to seedling disease burden (Figure 4A-D; Tables S4-S7). In the case of plant productivity, species-specific differences are attributed to mycorrhizal dependency associated with plant traits, including specific root length and specific leaf area (Romero et al., 2023). We hypothesize that species-specific differences in host competence under AMF inoculation might similarly depend on host traits. For example, plants displaying complex root systems (e.g., finely branched root systems) tend to be more susceptible to pathogen infection, yet mycorrhizal symbiosis can reduce infection in these plants, dependent upon colonization with a compatible mutualistic AMF partner (Sikes et al., 2009). Importantly, species-specific differences in host competence under AMF inoculation suggest a strong potential for AMF to reduce dilution effects, as the most competent hosts in the presence of AMF (i.e., disease amplifiers) may have low competence in the absence of AMF (i.e., act as disease diluters).

Experimental loss of AMF disrupted the ability to predict pathogen transmission from host community competence, but this effect was restricted to a single experimental species pool (Figure 3B, Table S9). This result underlies the need to consider diverse species assemblages in biodiversity-disease research, as strong interactions could result in seemingly idiosyncratic results if applied to the wrong groups of host species or in the wrong ecological contexts. Nevertheless, consistent with the prediction that AMF could modify dilution effects, our results revealed that host community competence only predicted disease risk in the presence of AMF (Figure 3A, Table S8). In other words, this experiment demonstrated, for the first time, that loss of a belowground mutualist can disrupt community regulation of disease risk. This result therefore extends past work suggesting that the presence of AMF is needed to maintain natural ecosystem functions, including relationships between plant biodiversity, productivity, and stability (Van Der Heijden et al., 1998).

Broadly, this study demonstrates that mutualists can increase disease risk in different plant communities, but that mutualistic relationships may also be necessary for critical host community regulation of disease transmission. These results were made evident through the implementation of a novel analytical approach that allowed us to quantify host species competence across biodiversity gradients over the course of ongoing epidemics. As a large component of threatened global diversity includes mutualistic interactions among different species (Bascompte & Jordano, 2007), these results underlie an increasing need to understand how ecosystems change across trophic levels. To effectively restore biodiversity and mitigate disease risk, researchers need to extend their focus beyond species richness alone and better integrate other species interactions, including mutualisms and host competence, as they could potentially emerge as crucial factors in forecasting disease risk in our changing world.

## Acknowledgements

We are grateful to M-L Spalinger, N. Zurbuchen, L. Mommer, E. Ampt, A-L. Laine, and to members of the Laine, Niklaus, and Disease Ecology and Diversity Labs for helpful comments and suggestions during the preparation of this work, as well as greenhouse and lab assistance from L. Gurtner, R. Teuber, M. Soares, M. McLaughlin, T. Voortman and M. Furler. This work was supported by the University of Zürich, Oregon State University, and an Ambizione Grant (PZ00P3_202027) from the Swiss National Science Foundation awarded to FWH.

## Author contributions

FWH & SEW conceptualized the project; AK, FWH, & PAN designed the study; AK & FWH performed the experiments and collected the data; AK & PAN analyzed the data; NAL, AK, & FWH wrote the first draft. All authors contributed substantially to revising the manuscript.

## Data availability

Upon acceptance, the data and code will be made available in a public repository such as Figshare. Data and code are available to reviewers as a zip-file uploaded with this manuscript

