## Appendix I: Supplemental Information for "Loss of a belowground mutualist disrupts a fundamental biodiversity-disease relationship"

Supporting Information

Appendix I – Supplemental Methods

Propagation and maintenance of the soil-borne root pathogen. We obtained an isolate of *Rhizoctonia solani* (AG4-HGI, strain CBS 124594) from the Westerdijk Fungal Biodiversity Institute in the Netherlands, following Ampt et al. (2022). The fungal pathogen was propagated from agar slants onto full-strength potato dextrose agar (PDA) and then maintained on 1/5 PDA (15 g agar-agar (Fischer Bioreagents, USA) and 7.8 g PDA (Sigma Chemical Co., USA) per liter of media) to sustain its virulence. To further sustain virulence, *R. solani* was only transferred twice to a fresh 1/5 PDA Petri dish, once every 21 days and once six days before the inoculation date of the experiment. The cultures were kept at 23 °C for growth and at 20 °C for storage in a dark climate chamber.

Substrate preparation and mycorrhizal inoculation. The substrate employed was a soil-sand mixture in a 2:1 ratio by weight. The soil (ED73 lawn soil (Einheitserde, Germany)) underwent sterilization by high-energy X-rays (25-60 kGy,). The quartz sand component was heat sterilized through autoclaving at 121 °C on two occasions. The substrate was reinoculated with a bacterial/rhizobia mix from an extensively managed grass meadow to establish a similar microbial community that would normally occur within the soil environment of the used species, following Weber et al. (2024). Specifically, 1 kg of soil substrate from an extensive meadow was mixed with 1.5 L of tap water, then filtered 2x through an 11 µm pore filter paper (MN 615, Macherey Nagel, Germany) to exclude mycorrhizal spores. The microbial wash derived from the aqueous suspension was diluted to 4 L with 1x filtered tap water. The mixture was added to the four trays of substrate while continuously mixing. Then 100 × 3 L round pots were packed for the AMF control treatment. For the mycorrhizal inoculum, we used a dosage of 100 g/mL * 300 L soil = 30 g of spores. The inoculum used was a mycorrhizal root powder (>381 million propagules per kg) preparation of intraradical spores and vesicles of the arbuscular mycorrhizal fungus *Rhizoglomus irregulare* (syn. *Rhizophagus irregularis,* strain QS81) from a muskmelon (Cucumis melo) batch (INOQ Gmbh, Germany). 7.5 g of spores were evenly added to each of the four trays and then evenly mixed again. Then the next 100 pots were packed for the AMF inoculation treatment and set aside.

Seed preparation and germination. To remove microbial contamination, the seeds were surfaced sterilized by 2% bleach for 20 minutes (seeds scarified with sandpaper in advance for Trifolium pratense) and then rinsed three times with demineralized water ($H_2O$). Seeds were then placed on the surface of Petri dishes filled with moisturized, heat-sterilized sand, covered with transparent covers. Seeds germinated after 2-3 days and were planted after five days. The seeds were obtained from a native seed supplier (UFA Samen, Switzerland).

Plant biodiversity gradient and species pools. As experiments are often carried out with a single replicate community, but understanding generality requires making comparisons across replicate communities, we decided to cluster the eight plant species into two different four-species pools. This grouping allows us to draw comparisons not only within individual communities but also across the different replicate communities. Each of the two-species pools contained one grass, one non-aster dicot, one aster, and one legume, with the identity of each species in each pool based on host competence. The biodiversity gradient was then established in each replicate community by planting four plants opposite one another in a random order at the edges of a square in the pot (Figure 1B). For the three-species communities, each replicate had a different duplicated species, so that duplicated species were evenly distributed across replicates. After one week, any missing seedlings (n = 41) were replaced with previously germinated seedlings. The plant communities were then grown for three weeks to support the establishment of a mycorrhizal network.

Planting of focal seedlings. Three weeks after the plant community was established, a row of seedlings of one plant species per species pool was planted into each community to serve as focal seedlings for measuring disease transmission. A previous attempt to plant all species in a grid into the center of the communities was unsuccessful, so we ultimately decided to choose only one focal species per pool to be planted in a row into the center of each pot. We selected two Asteraceae species (*Achillea millefolium* in pool one and *Leucanthemum vulgare* in pool two) to serve as focal species, because they showed intermediate competence in the pilot experiments and were phylogenetically matched. Focal seedlings were planted in a row of five individuals in the middle of each plant community, spaced two centimeters apart. During planting of the experiment, some seedling trays had lower germination, higher transplanting mortality, and lower establishment in the pots than expected. We therefore replaced dead seedlings with a backup batch or directly planted seeds to ensure timely planting in the experiment.

Experimental conditions in the greenhouse. To enable as constant abiotic conditions as possible, the experiment was conducted in controlled greenhouse conditions with a constant temperature of 24 °C ± 2 °C, 16 hours of light and 8 hours of darkness, light intensity of 180 µmol m-2 s-L and 40% relative humidity. During the experiment, the plants were misted from the top with a spray hose with demineralized water every day and watered from the bottom up in sterile bowls with the same amount of demineralized water once at the beginning and once in the middle of the experiment. The greenhouse tables with the pots were randomly arranged and rearranged three times to enable more equal abiotic conditions inside the greenhouse compartment. In the first 10 days, the pots were placed under transparent covers to allow *R. solani* to establish. The transparent covers were removed when the plants reached a height at which they could no longer support the covers.

Host competence measurements. To determine the susceptibility and transmissibility (i.e., competence) of each plant species to *Rhizoctonia solani*, we designed pilot experiments that were performed two months prior to the full experiment. The experiments were constructed as a linear transmission experiment where 5 seedlings of each species were placed at a two-centimeter distance in 500 mL pots filled with different substrates displaying varying soil conditions. All three pilot experiments were performed in controlled greenhouse conditions with a constant temperature of 24 °C ± 2 °C, 16 hours of light and 8 hours of darkness, light intensity of 180 µmol m-2 s-L, and 40 % relative humidity. During the experiment, the plants were bottom-watered in bowls with tap water every day and sprayed from the top at the beginning of each experiment to enable the seedlings to settle in the soil. The pots were arranged randomly in the greenhouse compartment, and transparent covers were used to increase humidity and aid in pathogen establishment. The seeds used were from a native seed supplier, UFA Samen. After establishment, the seedling at the leftmost position of each row was inoculated with *Rhizoctonia solani* by using an agar plug with actively growing mycelium (1 $cm^2$) from a 1/5 PDA culture plate (15 g agar-agar (Fischer Bioreagents, USA) and 7.8 g PDA (Sigma Chemical Co., USA) per liter of media). This was done by placing the agar plug with the mycelium side directly onto the primary root of the seedling and then loosely covering it with soil. From then on, the spread of disease was monitored from one infected individual to its neighboring seedlings every 1-3 days. Disease symptoms were measured as aboveground symptoms such as leaf and hypocotyl rot, discoloring or spotting, wilting, and stem collapse. At the end of the experiment, the number of diseased seedlings out of the five seedlings was recorded per plant species monoculture pot (Table S2).

For the first pilot experiment, we used non-sterilized standard lawn soil only. We planted 10 seeds directly into the soil, two in each hole with a dibble. The grass species seedlings were planted one day prior to the rest, and seeds of *Trifolium pratense* were scarified with sandpaper in advance. After five days, the seedlings in the pot were thinned out so that each pot only contained five established seedlings. Then the leftmost seedling was inoculated with *R. solani*. This experiment ran for 14 days.

The second and third pilot experiments more closely match the full experiment. First, seeds were sterilized using 2 % bleach for 20 minutes (seeds scarified with sandpaper in advance for *Trifolium pratense*) and rinsed with demineralized water three times. The seeds were then placed on the surface of Petri dishes filled with the moisturized, heat-sterilized sand, covered with transparent covers, and stored in the greenhouse compartment. After three days, seedlings were transplanted in a line of five into the substrate-filled pots. For the substrate, we used a 50:50 sand-to-soil ratio mixture by weight. The ED73 lawn soil used was sterilized by gamma X-ray (25- 60 kGy), and the quartz sand was sterilized through autoclaving at 121°C twice. The substrate was reinoculated with a bacterial/rhizobia mix by taking 500 g of soil substrate from an extensive meadow on campus for the total employed 2400 g of soil. The mass was mixed with 400 mL of tap water, and then filtered two times through 11 µm pore filter paper (MN 615, Macherey Nagel, Germany) to exclude mycorrhizal spores. Then the substrate was split in two and a first batch of pots was filled for the second pilot experiment. For the third pilot experiment, we added arbuscular mycorrhizal inoculum, with a dosage of 100 g/mL * 6 L soil = 600 mg. The mycorrhizal root powder (>381 million propagules per kg) preparation of intraradical spores and vesicles of the arbuscular mycorrhizal fungus *Rhizoglomus irregulare* from a muskmelon (Cucumis melo) batch (INOQ Gmbh, Germany) was evenly added to the substrate tray and mixed. In both experiments, the leftmost seedling was inoculated with *R. solani*. Both the second and third pilot experiments ran for seven days.

The values for "species competence" were summarized as the number of total infected plants per species over the three pilot experiments for both species pools and later used in the full experiment (Table S1). This measure captures both infection success of the pathogen as well as transmission among seedlings under different conditions.

Appendix II – Supplementary Tables and Figures

Table S1: Results from the Pilot Experiments

| Species | Abbreviation | Pilot 1 (Lawn soil) | Pilot 2 (+AMF) | Pilot 3 (-AMF) | Competence Score | Pool |
|---|---|---|---|---|---|---|
| Achillea millefolium | Am | 2 | 2 | 1 | 5 | 1 |
| Lotus corniculatus | Lc | 4 | 1 | 1 | 6 | 1 |
| Lolium perenne | Lp | 0 | 0 | 0 | 0 | 1 |
| Prunella vulgaris | Pv | 2 | 3 | 1 | 6 | 1 |
| Holcus lanatus | Hl | 1 | 1 | 0 | 2 | 2 |
| Leucanthemum vulgare | Lv | 1 | 2 | 2 | 5 | 2 |
| Plantago lanceolata | Pl | 1 | 2 | 1 | 4 | 2 |
| Trifolium pratense | Tp | 1 | 1 | 1 | 3 | 2 |
| Species Pool 1 | P1 | 8 | 6 | 3 | 17 | 1 |
| Species Pool 2 | P1 | 4 | 6 | 4 | 14 | 2 |

Table S2: Type I Analysis of Variance Table for the interactive effects of planted diversity and mycorrhizal inoculation on disease burden

| | Sum Sq | Mean Sq | NumDF | DenDF | F-value | P-value |
|---|---|---|---|---|---|---|
| **Diversity** | 177.96 | 177.96 | 1 | 28 | 2.332 | 0.138 |
| **Mycorrhiza** | 330.55 | 330.55 | 1 | 28 | 4.332 | 0.047 * |
| **Diversity x Mycorrhiza** | 30.59 | 30.59 | 1 | 28 | 0.401 | 0.532 |

Table S3: Type I Analysis of Variance Table for the interactive effects of planted diversity, mycorrhizal inoculation, and plant species pool on disease burden

| | Sum Sq | Mean Sq | NumDF | DenDF | F-value | P-value |
|---|---|---|---|---|---|---|
| **Diversity** | 148.13 | 148.13 | 1 | 26 | 3.404 | 0.076 . |
| **Mycorrhiza** | 330.55 | 330.55 | 1 | 26 | 7.560 | 0.0105 * |
| **Pool** | 619.35 | 619.35 | 1 | 26 | 14.232 | <0.001 *** |
| **Diversity x Mycorrhiza** | 30.59 | 30.59 | 1 | 26 | 0.703 | 0.409 |
| **Diversity x Pool** | 27.54 | 27.54 | 1 | 26 | 0.633 | 0.433 |
| **Mycorrhiza x Pool** | 983.94 | 983.94 | 1 | 26 | 22.610 | <0.001 *** |
| **Diversity x Mycorrhiza x Pool** | 21.09 | 21.09 | 1 | 26 | 0.485 | 0.492 |

Table S4: Diallel Analysis for Species Pool 1, -AMF, see Table S1 for species abbreviations

| | GCA Estimate | Std. Error | t-value | P-value |
|---|---|---|---|---|
| **Am** | 16.19 | 2.38 | 6.806 | <0.001 *** |
| **Pv** | 16.67 | 2.38 | 7.009 | <0.001*** |
| **Lc** | 6.66 | 2.38 | 2.801 | 0.0311* |
| **Lp** | 10.54 | 2.38 | 4.431 | 0.004 ** |

**Residual standard error: 6.142 on 6 degrees of freedom**

**Multiple R-squared: 0.967, Adjusted R-squared: 0.945**

**F-statistic: 44.27 on 4 and 6 DF, p-value: <0.001**

Table S5: Diallel Analysis for Species Pool 1, +AMF, see Table S1 for species abbreviations

| | GCA Estimate | Std. Error | t-value | P-value |
|---|---|---|---|---|
| **Am** | 24.20 | 4.47 | 5.413 | 0.002** |
| **Pv** | 27.32 | 4.47 | 6.113 | <0.001*** |
| **Lc** | 16.08 | 4.47 | 3.596 | 0.011* |
| **Lp** | 12.49 | 4.47 | 2.795 | 0.031* |

**Residual standard error: 11.540 on 6 degrees of freedom**

**Multiple R-squared: 0.955, Adjusted R-squared: 0.925**

**F-statistic: 31.71 on 4 and 6 DF, p-value: <0.001**

Table S6: Diallel Analysis for Species Pool 2, -AMF, see Table S1 for species abbreviations

| | GCA Estimate | Std. Error | t-value | P-value |
|---|---|---|---|---|
| **Lv** | 7.35 | 1.863 | 3.948 | 0.008** |
| **Pl** | 16.50 | 1.863 | 8.858 | <0.001*** |
| **Tp** | 15.70 | 1.863 | 8.426 | <0.001*** |
| **Hl** | 11.11 | 1.863 | 5.963 | <0.001*** |

**Residual standard error: 4.809 on 6 degrees of freedom**

**Multiple R-squared: 0.980, Adjusted R-squared: 0.966**

**F-statistic: 72.87 on 4 and 6 DF, p-value: <0.001**

Table S7: Diallel Analysis for Species Pool 2, +AMF, see Table S1 for species abbreviations

| | GCA Estimate | Std. Error | t-value | P-value |
|---|---|---|---|---|
| **Lv** | 10.07 | 2.50 | 4.026 | 0.007** |
| **Pl** | 13.93 | 2.50 | 5.569 | 0.001** |
| **Tp** | 10.42 | 2.50 | 4.167 | 0.006** |
| **Hl** | 10.05 | 2.50 | 4.019 | 0.007** |

**Residual standard error: 6.456 on 6 degrees of freedom**

**Multiple R-squared: 0.952, Adjusted R-squared: 0.921**

**F-statistic: 30.02 on 4 and 6 DF, p-value: <0.001**

Table S8: Type I Analysis of Variance Table for model testing interactive effects of host community competence and mycorrhizal inoculation on disease burden

| | Sum Sq | Mean Sq | NumDF | DenDF | F-value | P-value |
|---|---|---|---|---|---|---|
| **Host community competence (pilot data)** | 277.37 | 277.37 | 1 | 28 | 4.535 | 0.042 * |
| **Mycorrhiza** | 330.55 | 330.55 | 1 | 28 | 5.404 | 0.028 * |
| **Host community competence x Mycorrhiza** | 454.47 | 454.47 | 1 | 28 | 7.430 | 0.011 * |

Table S9: Type I Analysis of Variance Table for model testing interactive effects of host community competence, mycorrhizal inoculation, and host species pool on disease burden

| | Sum Sq | Mean Sq | NumDF | DenDF | F-value | P-value |
|---|---|---|---|---|---|---|
| **Host community competence (pilot data)** | 229.22 | 229.22 | 1 | 26 | 5.866 | 0.023 * |
| **Pool** | 379.94 | 379.94 | 1 | 26 | 9.723 | 0.004 ** |
| **Mycorrhiza** | 330.55 | 330.55 | 1 | 26 | 8.459 | 0.007 ** |
| **Host community competence x Pool** | 19.38 | 19.38 | 1 | 26 | 0.496 | 0.488 |
| **Host community competence x Mycorrhiza** | 454.47 | 454.47 | 1 | 26 | 11.631 | 0.002 ** |
| **Pool x Mycorrhiza** | 696.66 | 696.66 | 1 | 26 | 17.829 | <0.001 *** |

| | | | | | | |
|---|---|---|---|---|---|---|
| **Host community competence x Pool x Mycorrhiza** | 0.00 | 0.00 | 1 | 26 | 0.0000 | 0.995 |

Figure S1-S8: Microscopy photos of arbuscular mycorrhizal fungi (AMF) colonization

| | |
|---|---|
| 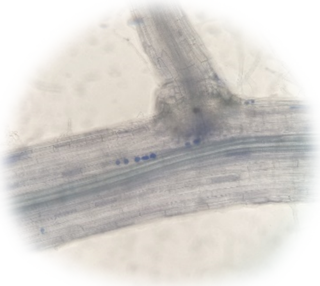 | 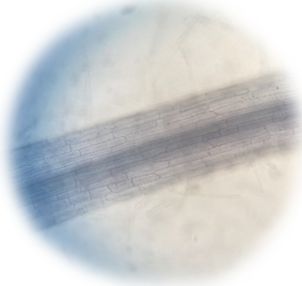 |
| Figure S1A: Pot Nr. 1 (+AMF), Support plant 4, Achillea millefolium, Light microscope 200x | Figure S1B: Pot Nr. 2 (-AMF), Support plant 1, Achillea millefolium, Light microscope 200x |
| 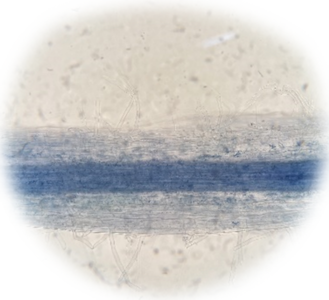 | 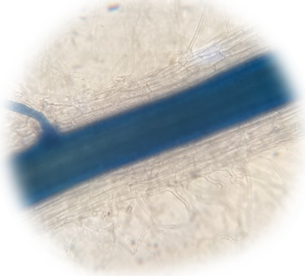 |
| Figure S2A: Pot Nr. 9 (+AMF), Support plant 2, Holcus lanatus, Light microscope 200x | Figure S2B: Pot Nr. 10 (-AMF), Support plant 1, Holcus lanatus, Light microscope 200x |
| 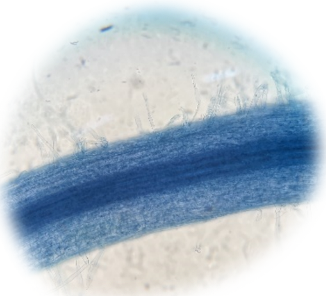 | 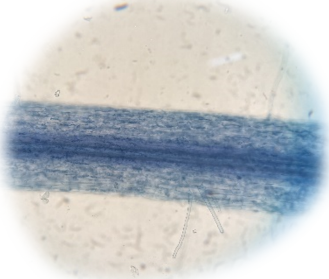 |
| Figure S3A: Pot Nr. 17 (+AMF), Support plant 2, Lotus corniculatus, Light microscope 200x | Figure S3A: Pot Nr. 18 (-AMF), Support plant 3, Lotus corniculatus, Light microscope 200x |

| | |
|---|---|
| 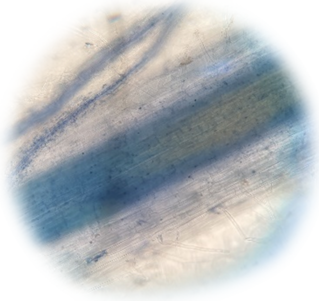 | 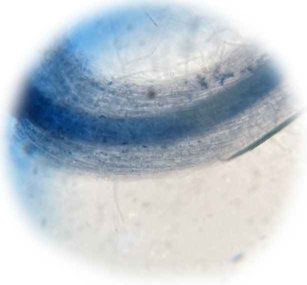 |
| Figure S4A: Pot Nr. 25 (+AMF), Support plant 4, Lolium perenne, Light microscope 200x | Figure S4A: Pot Nr. 26 (-AMF), Support plant 1, Lolium perenne, Light microscope 200x |
| 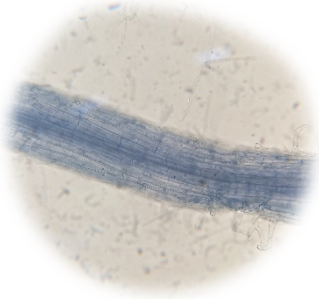 | 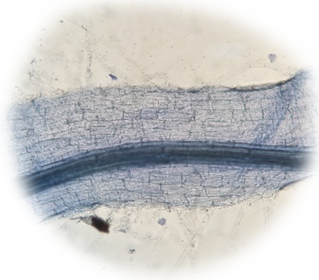 |
| Figure S5A: Pot Nr. 33 (+AMF), Support plant 3, Leucanthemum vulgare, Light microscope 200x | Figure S5B: Pot Nr. 34 (-AMF), Support plant 4, Leucanthemum vulgare, Light microscope 200x |
| 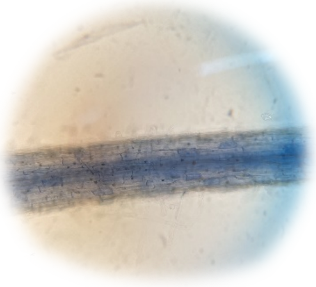 | 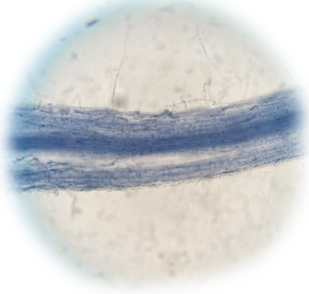 |
| Figure S6A: Pot Nr. 41 (+AMF), Support plant 1, Plantago lanceolata, Light microscope 200x | Figure S6B: Pot Nr. 42 (-AMF), Support plant 4, Plantago lanceolata, Light microscope 200x |

| | |
|---|---|
| 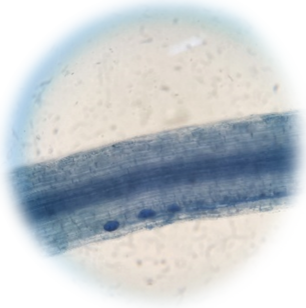 | 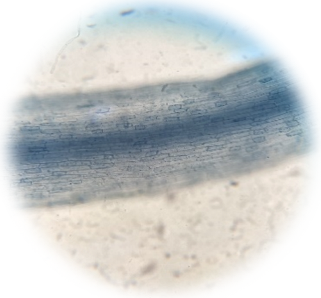 |
| Figure S7A: Pot Nr. 49 (+AMF), Support plant 3, Prunella vulgaris, Light microscope 200x | Figure S7B: Pot Nr. 50 (-AMF), Support plant 2, Prunella vulgaris, Light microscope 200x |
| 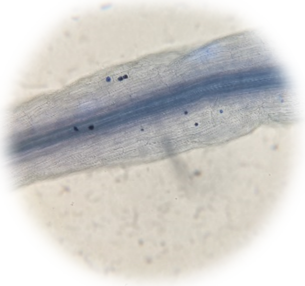 | 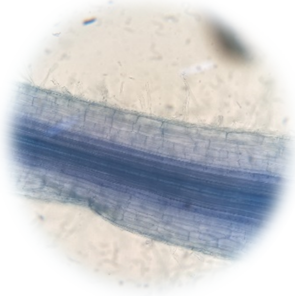 |
| Figure S8A: Pot Nr. 57 (+AMF), Support plant 1, Trifolium pratense, Light microscope 200x | Figure S8B: Pot Nr. 58 (-AMF), Support plant 1, Trifolium pratense, Light microscope 200x |
